# Loss of cell cycle gatekeeping by CNOT3 impairs hematopoietic stem and progenitor cell division and repopulating activity

**DOI:** 10.1101/2025.10.23.684176

**Authors:** Katie Yuen, Thilelli Taibi, Quang Anh Hoang, Marty Yue, Zongmin Liu, Effat Habibi, Elias Spiliotopoulos, Niveditha Ramkumar, Nick Nation, Glenn Edin, Thuy T M Ngo, Ly P. Vu

## Abstract

Adult mammalian hematopoietic stem cells (HSCs) constitute a heterogeneous population responsible for generating various cell types in the blood throughout adulthood. Gene expression programs underlying regulation of self-renewal and differentiation of HSCs are tightly regulated. However, how post-transcriptional regulation of gene expression influences HSCs and hematopoiesis remains largely unexplored. Here, we report the critical role of CNOT3, a subunit of the CCR4-NOT complex, in regulating hematopoietic stem cells (HSCs) function in adult hematopoiesis. We observed that *Cnot3* mRNA is highly expressed in HSCs and CNOT3 ablation in the murine *Cnot3* conditional knockout mouse model resulted in anemia, reduced bone marrow cellularity and enhanced extramedullary hematopoiesis in spleen. Deletion of *Cnot3* resulted in the early expansion of immunophenotypic HSCs which were then progressively lost over time. *Cnot3* knockout hematopoietic stem/progenitor cells (HSPCs) failed to reconstitute hematopoietic systems of recipient animals in transplantation assays. Single-cell RNA sequencing (scRNA-seq) analysis of HSPCs revealed disruptions in lineage development and loss of HSCs. Transcriptomic profiling and cell cycle analysis demonstrated that *Cnot3* deletion led to increased cycling activity in HSCs. Our results indicate that CNOT3 is critical for maintenance of homeostasis in HSCs and the hematopoietic system.

## Introduction

Hematopoietic stem and progenitor cells (HSPCs) are responsible for the lifelong production of all blood lineages, a process requiring a delicate balance between stem cell self-renewal and multi-lineage differentiation. These fate decisions demand precise coordination of gene expression programs to ensure that sufficient mature blood cells are produced while the stem cell pool is maintained. It is well established that a coordinated network of master transcription factors and epigenetic modifiers plays a central role in orchestrating these delicate processes during hematopoiesis(1, 2). On the other hand, post-transcriptional regulation has increasingly been recognized as an equally fundamental layer of gene expression control. Post-transcriptional mechanisms refer to a set of diverse and complex processes governing the entire life cycle of an mRNA molecule from mRNA splicing, capping, 3’ end formation to nucleocytoplasmic export to the cytoplasm before being translated by ribosomes and then degraded(3). Post-transcriptional regulation could specifically target key transcription factors, cell cycle regulators, and signaling effectors to mediate key biological responses(4, 5). A number of studies had illustrated critical requirements of RNA regulating proteins such as Lin28b(6), MSI2(7, 8), SYNCRIP(9) and RNA methyltransferase METTL3(10) in hematopoietic stem cells’ function and blood formation, underscoring the importance of understanding how these mechanisms contribute to maintenance of stem cells and normal development(11–13). At the same time, they also called for expanded efforts to further elucidate the relevance and requirement of post-transcriptional gene expression control in hematopoietic stem cells (HSCs) and hematopoiesis.

The evolutionarily conserved CCR4-NOT (CNOT) complex is a multi-subunit machinery which participates in multiple layers of gene expression control i.e., transcription, mRNA decay and translation(14). Through these activities, the CNOT complex plays a key role in maintaining proper gene expression homeostasis in many biological contexts(15). Among CNOT components, the regulatory subunit CNOT3 while lacking intrinsic enzymatic activity can stabilize the complex and recruit other factors to specific mRNA targets. Emerging evidence indicates that CNOT3 is critical for maintenance of cell proliferation and differentiation in a context-dependent manner. In pluripotent embryonic stem cells (ESCs), CNOT3 (together with CNOT1 and CNOT2) is required to maintain the undifferentiated state, as depletion of these subunits causes loss of stem cell identity and premature differentiation into extraembryonic lineages(16, 17). CNOT3 is required to promote degradation of differentiation genes to maintain ESCs’ pluripotency(17). Conversely, in lineage-committed progenitors such as developing B lymphocytes, CNOT3 is required to enable differentiation and proliferation. Conditional knockout of *Cnot3* in B cells blocks the pro- to pre-B cell transition due to a failure in immunoglobulin heavy chain gene rearrangement and aberrant stabilization of *Trp53* (*p53*) mRNA(18). In addition, loss of one *Cnot3* allele in Cnot3-heterozygous mice results in aberrant energy metabolism(19), decreased bone formation(20), defects in heart(21) and adipocyte function(20). These examples underscore how CNOT3 exerts precise post-transcriptional control over key transcripts to influence diverse physiological processes. CNOT3 has also been implicated in disease settings. CNOT3 was identified as a recurrently mutated tumor suppressor in T-cell acute lymphoblastic leukemia (T-ALL), with loss-of-function mutations found in ∼8% of cases(20). In contrast, other cancers appear to co-opt CNOT3 for tumor-promoting effects. High CNOT3 expression is associated with more advanced, poor-prognosis colon carcinoma(22) and upregulation of CNOT3 promotes development of non-small cell lung cancer by suppressing expression of KLF2 and p21(23). Our recent work showed that CNOT3 is critical for sustaining the growth of myeloid leukemia cells(24). As these findings collectively point to CNOT3 as a pivotal regulator of cell fate, it is important to define CNOT3’s exact role and requirement in various developmental or oncogenic contexts.

In this study, we developed a conditional and tissue-specific genetic deletion model in mice to examine the function of CNOT3 in adult hematopoiesis. The multilineage nature of the hematopoietic system and the exquisite self-renewal activity of the HSPCs provides a unique opportunity to dissect the functional requirement and the gene expression programs regulated by CNOT3 in different blood cell types simultaneously. Our extensive phenotypic characterization and single cell analysis revealed lineage specific impacts of CNOT3 depletion and its requirement for HSPCs function. Cell cycle and division analyses revealed that loss of CNOT3 drives HSCs to exit quiescence and enter rapid proliferation, but fail to sustain long term division and self-renewal. Overall, our study uncovered CNOT3 as a critical regulator responsible for restraining HSPCs from cycling, hence plays an important role in maintenance of HSPCs.

## Results

### *Cnot3* deletion resulted in anemia, loss of hematopoietic stem cells and hematopoietic failure

We surveyed the expression of *Cnot3* in the hematopoietic system and observed that *Cnot3* is highly expressed in hematopoietic stem/progenitor cells (HSPCs) and downregulated in differentiated cells (**Figure S1A-B**). To explore the role of CNOT3 in HSC and hematopoiesis, we crossed *Cnot3^flox/flox^*(*17*) (**Figure S1C**) with interferon (IFN)-inducible *Mx1-Cre+* transgenic mice to generate *Cnot3^flox/flox^/*Cre+ mice. Cre can be activated by administration of a IFN inducer polyInosinic-polyCytidylic acid (pIpC). We injected *Cnot3^flox/flox^/*Cre- and *Cnot3^flox/flox^/*Cre+ mice with pIpC for 2 consecutive days and at 3 weeks post-pIpC, effective deletion of the targeted region was confirmed by PCR (**Figure 1A-B**). Ablation of protein expression was demonstrated by immunoblots (**Figure S1D-E**). After pIpC treatment, *Cnot3^flox/flox^/*Cre- and *Cnot3^flox/flox^/*Cre+ animals are referred to as control and cKO mice respectively.

**Figure 1.**
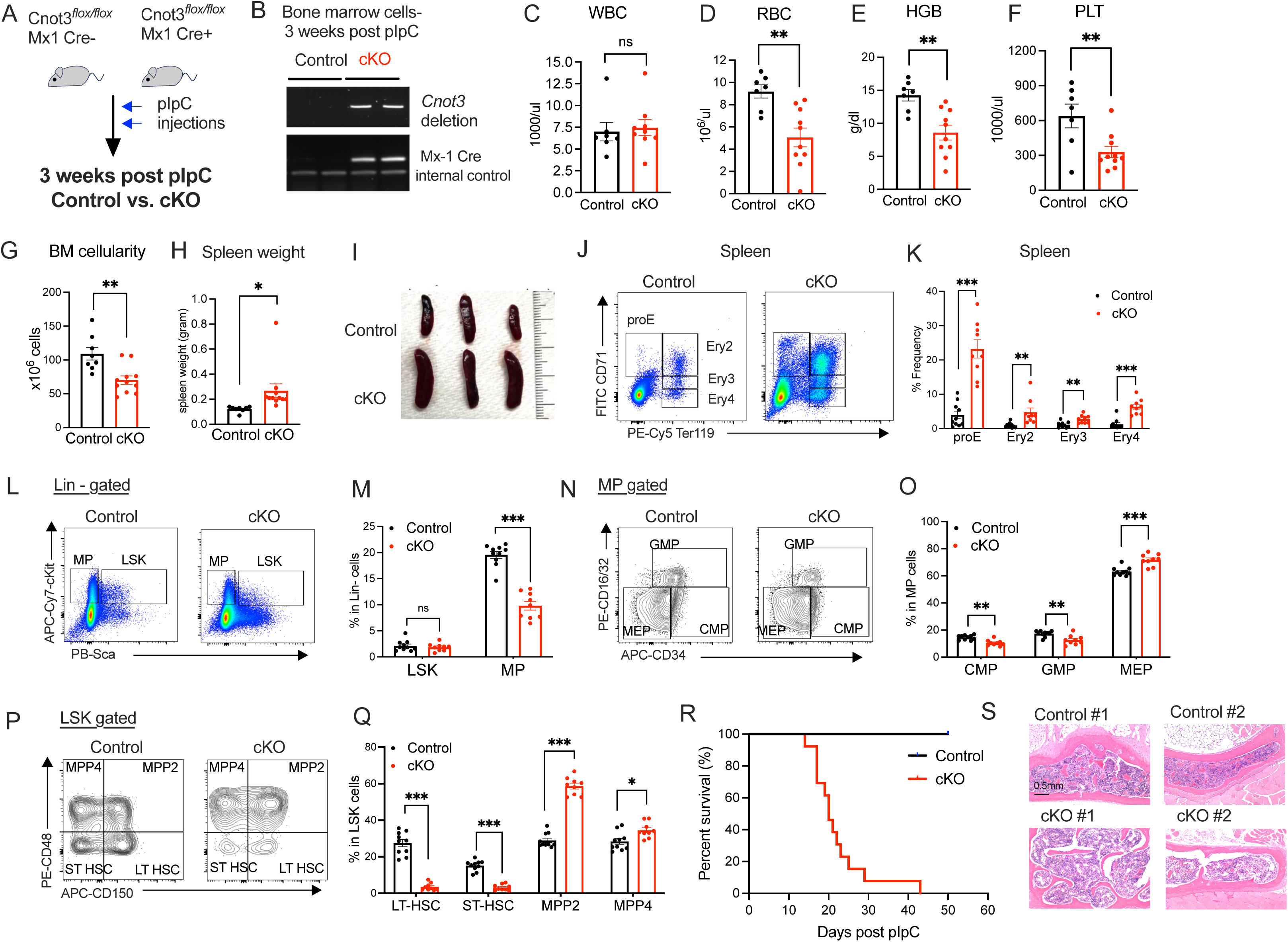
*Cnot3* deletion resulted in hematopoietic failure and depletion of HSCs (A) Experimental scheme for pIpC injection and analysis of hematopoietic system upon deletion of *Cnot3* at 3 weeks post-pIpC. (B) Effective deletion of *Cnot3* demonstrated by PCR by using genomic DNA of bone marrow cells from pIpC treated Control vs. cKO mice. Primers used were indicated in Figure S1C. (C-F) Analysis of blood counts: (C) White blood count (WBC); (D) Red Blood count (RBC); (E) Hemoglobin (HGB) and (F) Platelets (PLT) of Control vs. cKO mice at 3 weeks post-pIpC. (G) Total bone marrow cellularity of Control vs. cKO mice. (H) Spleen weight of Control vs. cKO mice. (I) Representative images of spleen of Control vs. cKO mice. (J) Representative flow plots of erythroid cells from spleen of Control vs. cKO mice. ProE: Proerythroblasts; Ery2-4: Erythroid 2-4. (K) Frequencies of different erythroid subpopulations from spleen of Control vs. cKO mice. (L) Representative flow plots of gating strategy for LSK (Lin-cKit+ Sca-1+) and MP (Myeloid progenitor, Lin-cKit+ Sca-1-) cells in bone marrow (BM). (M) Frequencies of LSK and MP in Control vs. cKO mice. (N) Representative flow plots of gating strategy for CMP, GMP and MEP. (O) Frequencies of CMP, GMP and MEP in Control vs. cKO mice. (P) Representative flow plots of gating strategy for LT-HSC (LSK, CD150+CD48-), ST-HSC (LSK, CD150-CD48-), MPP2 (LSK, CD150+CD48+), and MPP4 (LSK, CD150-CD48+). (Q) Frequencies of LT-HSC, ST-HSC, MPP2 and MMP4 in Control vs. cKO mice. (R) Survival curve of Control (n=10) vs. cKO (n=13) mice after pIpC injection. Log-rank test was used. (S) Representative images of H&E staining of Sternal bone marrow samples from Control vs. cKO mice. All graphs show data as mean+/- s.e.m, n=7-10 in Control vs. n=9-10 in cKO. Two-tailed Student’s t test. ns not significant * p< 0.05, **p<0.01, *** p<0.001.

Complete blood count (CBC) analysis showed that while white blood cells were at a similar level in cKO vs. control, there were a marked decrease in counts of red blood cells (RBC) and platelets (PLT) as well as lower Hemoglobin (HGB) levels in cKO mice (**Figure 1C-F**). *Cnot3* cKO mice exhibited reduced bone marrow (BM) cellularity and enlarged spleen (**Figure 1G-I**). Multiparameter flow cytometry analysis of peripheral blood (PB), bone marrow (BM) and spleen revealed defects in erythroid development in *Cnot3* KO animals, which were characterized by the reduction of mature erythroid cells in both PB and BM (**Figure S1F-G**). In addition, we observed that while circulating myeloid cells including granulocytes and monocytes in PB were equally represented in control vs. cKO mice, there was a marked decrease in Gr1+ Mac1+ mature granulocyte in BM but not in PB (**Figure S1F-G)**. On the other hand, there were enhanced extramedullary hematopoiesis characterized by increased red blood cell production (**Figure 1J-K**) and accumulation of mature granulocyte at the expense of B and T cells in the spleen (**Figure S1H-I**). Taken together, the data indicated that CNOT3 loss of function led to multi-lineage defects, in particular defects in differentiation of erythroid and myeloid cells.

Next, we examined the hematopoietic stem and progenitor compartments upon CNOT3 depletion. We observed a significant drop in frequency of the myeloid progenitors (MP-Lin-Sca-ckit+) and a mild increase in MEP (Lin-Sca-ckit+ CD34lowCD16/32-) concurrently with a decrease in CMP (Lin-Sca-ckit+ CD34highCD16/32-) and GMP (Lin-Sca-ckit+ CD34highCD16/32+) (**Figure 1L-O**). While there was no change in the percentage of the LSK (Lin-Sca+ckit+) population, further immunophenotypic analysis of the HSPCs showed a drastic loss of long-term HSCs (LT-HSCs, LSK CD48-CD150+) and short-term HSCs (ST-HSCs; LSK CD48-CD150-) and expansion of the downstream progenitors MPP2 (LSK CD48+CD150+) and MPP4 (LSK CD48+CD150-) (**Figure 1P-Q**). During the course of up to 50-day observation, all *Cnot3*-cKO died with the median survival of 3 weeks post-pIpC (**Figure 1R**). Prior to the time of death, *Cnot3*-cKO mice exhibited pale paws and femur and CBC analysis showed anemia, reduced platelets but notably no reduction in total WBC (**Figure S1K-N**). Pathological analysis confirmed a notable increase in erythropoietic activity in the spleen (**Figure S1J)** and reduced hemopoietic activity in the BM where the cell population was predominantly neutrophilic series cells (**Figure 1S**). In animals succumbed to death including those died earlier than 3 weeks post-pIpC, we also observed a complete abolishment of the HSC populations and proportional gain of MPPs upon CNOT3 depletion (**Figure S1O)**. These data indicated that CNOT3 is critically important to maintain homeostasis of the HSPC compartments.

### CNOT3 is required for repopulating activity of HSPCs

As loss of immunophenotypic HSCs might not necessarily translate to functional defects, we performed bone marrow transplantation to test the repopulating capacity of *Cnot3* deficient cKO HSPCs (**Figure 2A**). 2×10^6^ total BM cells from donor mice (CD45.2; control vs. cKO) were injected into lethally irradiated recipient mice (CD45.1). Bone marrow (BM) was analyzed at 8, 16 and 24 weeks after transplantation and complementary peripheral blood (PB) analysis was performed at 6, 12 and 34 weeks after transplantation to track chimerism of donor cells. While the control donor cells engrafted effectively both within the BM and in PB, *Cnot3*-cKO cells failed to reconstitute the hematopoietic system of recipient animals (**Figure 2B-C and Figure S2A**). Donor derived chimerism levels were minimal across all mature lineages (**Figure S2B**) as well as the stem and progenitor compartments (**Figure 2D**).

**Figure 2.**
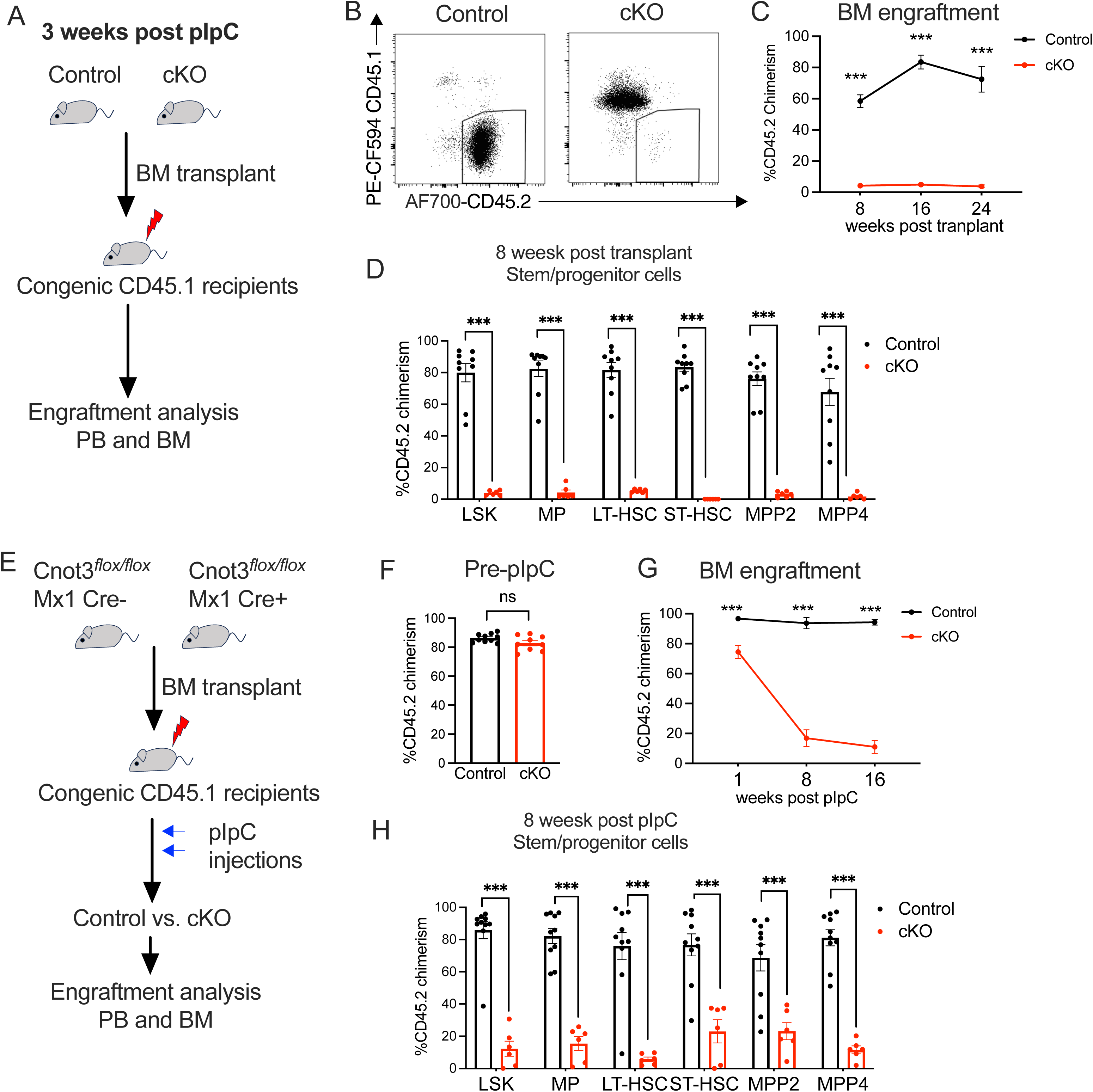
*Cnot3* deletion abolishes repopulating ability of HSPCs (A) Scheme for experimental strategy to transplant BM cells isolated from Control vs. cKO mice depicted in figure 1A. (B) Representative flow plots showing chimerism of Control vs. cKO engrafted donor (CD45.2) cells vs. recipient (CD45.1) cells in BM of recipient animals. (C). Quantitative analysis of donor-derived chimerism at 8-, 16-and 24-weeks post-transplantation in BM of recipient animals. (D) Donor chimerism of Control vs. cKO in BM stem/progenitor populations at 8 weeks post-transplantation. (E) Scheme showing transplant strategy and experimental procedures. Congenic recipient mice were transplanted with *Cnot3*^flox/flox^ Cre- or Cre+ BM cells (pre-pIpC). After 6 weeks post-transplantation when engraftment of donor cells reached a high and stable level, recipient mice were injected with pIpC to delete *Cnot3*. (F) Equal donor chimerism of Control vs. cKO in BM of recipient animals at pre-pIpC. (G) Quantitative analysis of donor-derived chimerism at 1-, 8-and 16-weeks post-pIpC in BM of recipient animals. (H) Donor chimerism of Control vs. cKO in BM stem/progenitor populations at 8 weeks post-transplantation. All graphs show data as mean+/- s.e.m, n=2 donors for each genotype; n=10 recipients Control vs. n= 6 recipients cKO. Two-tailed Student’s t test. ns not significant, *** p<0.001.

To exclude the possible impact of *Cnot3* deletion in non-hematopoietic cells such as stromal cells in donors prior to transplantation, we transplanted *Cnot3^flox/flox^/*Cre- and *Cnot3^flox/flox^/*Cre+ total BM cells and allowed donor cells to reach stable engraftment in recipient animals before administration of pIpC (**Figure 2E-F**). Upon induction of Cre, we observed a drop in donor chimerism in recipients engrafted with *Cnot3*-cKO cells at 1-week post-pIpC and engraftment of *Cnot3*-cKO cells was progressively abrogated at later time points of 8 and 16 weeks while chimerism in control recipient animals was not impacted (**Figure 2G**). *Cnot3*-cKO cells also displayed a significantly reduced engraftment in the HSPC compartments (**Figure 2H**). Taken together, these data strongly supported the cell-intrinsic role of CNOT3 in regulating HSPCs’ function.

### Depletion of CNOT3 resulted in functionally defective HSCs

Given the close to complete loss of HSC populations in *Cnot3*-cKO animals at 3 weeks post-pIpC, we sought to characterize the course of changes within the HSPC compartments and to capture the timepoint prior to abolition of HSCs upon CNOT3 KO. We performed comprehensive analysis of the hematopoietic system at 1-week and 2-week post induction of *Cnot3* deletion (**Figure 3A and S3G**). At 1 week post pIpC, we observed no major change in blood counts except a reduction in platelets of *Cnot3*-cKO mice (**Figure S3A-D**). There was a decrease in BM cellularity in some animals but no increase in spleen weight (**Figure S3E-F**). *Cnot3*-cKO mice displayed equivalent frequencies of LSKs and MPs in comparison to control (**Figure 3B**). Notably, in contrast to what observed at the 3 week-time point, both LT-HSC and ST-HSC populations were significantly expanded in the expense of MPP2 and MPP4 progenitors in *Cnot3*-cKO animals (**Figure 3C-D**). At the 2-week time point, the HSC expansion was no longer observed and instead HSPC frequencies more closely resembled those observed at 3 weeks post pIpC (**Figure S3G-H**). At the same time, defects in peripheral blood counts became apparent (**Figure S3I-L**). The data suggested that CNOT3 depletion resulted in initial expansion of HSC populations, which were then experienced gradual loss over time. As we observed the intact HSC populations at 1 week after *Cnot3* was deleted, we performed BM transplantation assay to determine whether they are functionally effective (**Figure 3E**). Interestingly, despite being immunophenotypically normal, *Cnot3*-cKO HSCs did not engraft in neither BM nor PB of recipient mice (**Figure 3F-G**). Almost no contribution of *Cnot3*-cKO donor cells were detected within HSPC compartments (**Figure 3H**). To further determine the dosage requirement for CNOT3 in sustaining repopulating activity, we generated *Cnot3*-cKO het, which lost one *Cnot3* allele (**Figure S3M**). Upon transplantation of bone marrow cells from Control vs. *Cnot3*-cKO het, we observed that reduction of CNOT3 by 50%, as demonstrated by immunoblot (**Figure S3N**) did not impact the engraftment of donor cells (**Figure S3O**). The results indicated that one allele of *Cnot3* is sufficient while there is a certain threshold required for CNOT3 activity to maintain HSPCs’ function.

**Figure 3.**
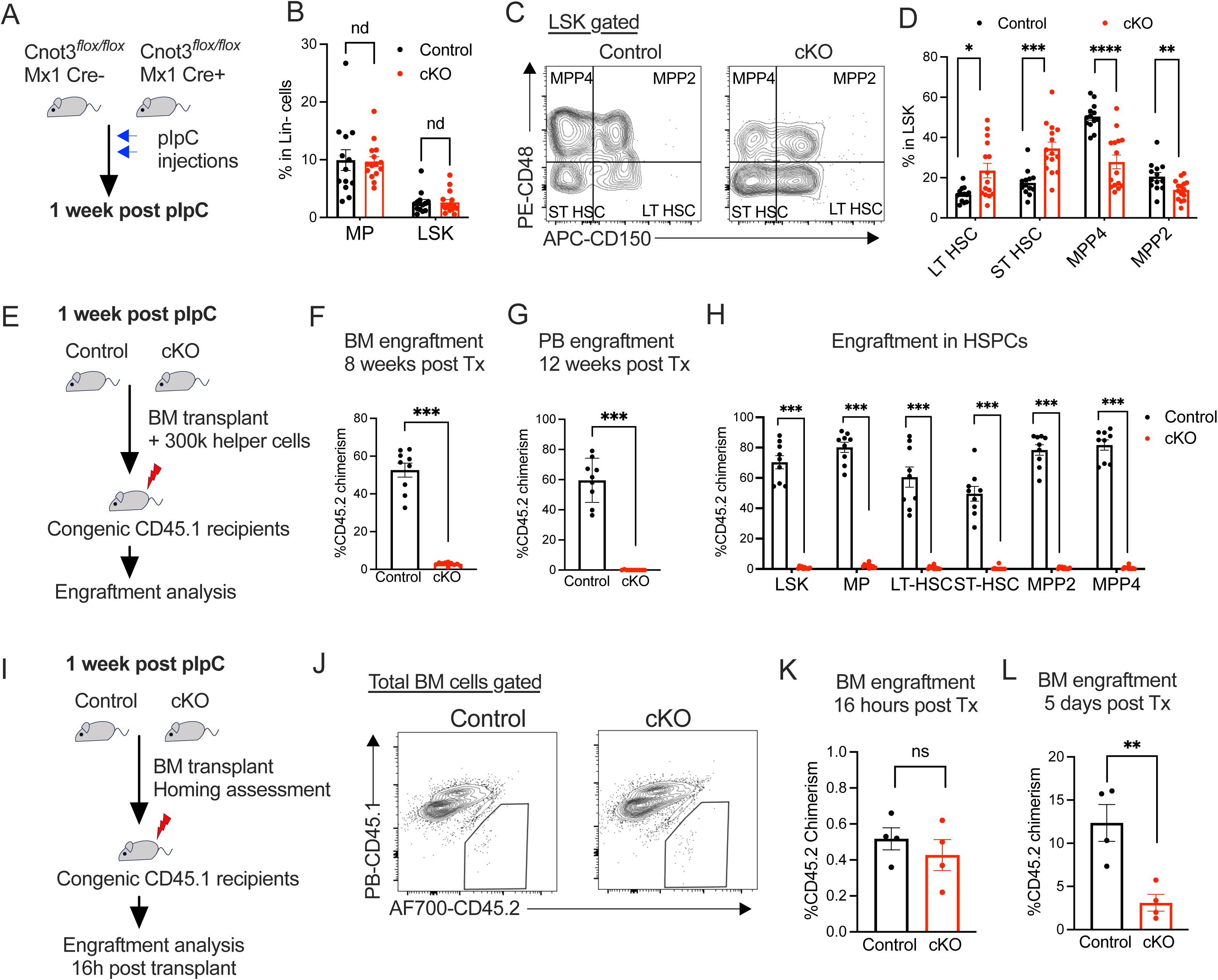
Normal Immunophenotypic *Cnot3*-cKO HSPCs are functionally defective (A) Experimental scheme for pIpC injection and analysis of hematopoietic system upon deletion of *Cnot3* at 1 weeks post-pIpC. (B) Frequencies of LSK and MP in Control vs. cKO mice. (C) Representative flow plots of gating strategy for LT-HSC (LSK, CD150+CD48-), ST-HSC (LSK, CD150-CD48-), MPP2 (LSK, CD150+CD48+), and MPP4 (LSK, CD150-CD48+). (D) Frequencies of LT-HSC, ST-HSC, MPP2 and MMP4 in Control vs. cKO mice. (E) Scheme for experimental strategy to transplant BM cells isolated from Control vs. cKO mice depicted in figure 3A. (F) Donor chimerism of Control vs. cKO in BM at 8 weeks post-transplantation. (G) Donor chimerism of Control vs. cKO in peripheral blood (PB) at 12 weeks post-transplantation. (H) Donor chimerism of Control vs. cKO in BM stem/progenitor populations at 8 weeks post-transplantation. (I) Scheme for experimental strategy to transplant BM cells isolated from Control vs. cKO mice depicted in figure 3A and followed up for bone marrow homing. (J) Representative flow plots showing detection of donor cells in bone marrow of recipient mice 16h post-transplant. (K-L) Donor chimerism of Control vs. cKO in BM at (K) 16 hours and (L) 5 days post-transplant. All graphs show data as mean+/- s.e.m. For primary cohorts, n=9-13 Control vs. n=10-15 cKO. For transplant, n=2 donors for each genotype; n=9 recipients Control vs. n= 9 recipients cKO. For homing assays, n=2 for each genotype; n=4 recipients for both Control and cKO. All graphs show data as mean+/- s.e.m. Two-tailed Student’s t test. ns no significant * p< 0.05, **p<0.01, *** p<0.001.

To directly interrogate the possibility that the engraftment defect might be due to impaired homing activity of HSPCs, we carried out the transplantation assay and examined the presence of donor cells from control vs. cKO mice in the BM of recipients 16 hours post-transplant (**Figure 3I**). We found that the *Cnot3*-cKO cells were able to effectively home to the BM similarly to the control cells (**Figure 3J-K**). At 5 days after transplantation, while increase in chimerism was observed in all recipients, *Cnot3*-cKO cells engrafted at a significantly lower level than that of control cells (**Figure 3L**). The data suggested that CNOT3 is not required for homing of HSPCs to the BM but is critically important for maintenance of HSCs function.

### Single cell transcriptomic profiling revealed multilineage-abnormalities upon CNOT3 depletion

To pinpoint the effect of *Cnot3* deletion on lineage commitment and cellular identity and uncover the molecular mechanisms for impaired function of *Cnot3*-cKO HSCs, we performed single-cell RNA-seq (scRNA-seq) analysis of HSPCs (Lin-cKit+) cells isolated from three paired control vs. cKO mice at 1-week post-pIpC (**Figure 4A**). Uniform Manifold Approximation and Projection (UMAP) analysis of combined control and cKO samples captured the majority of cell populations as previously described (10, (25) (**FigureS4A, Figure 4B and Table S1**). Upon *Cnot3* deletion, transcriptomic profiles of many cell clusters were extensively altered (**Figure 4C-D**). In agreement with the reported role of CNOT3 in B cell development, there was a complete loss of preB populations (**Figure 4E**). In addition, we observed a significant reduction of erythroid cells and a clear shift within the myeloid lineage (**Figure 4E**). Within the stem/progenitor compartments, CNOT3 depletion resulted in marked decrease in HSC frequency and noticeable changes in representation of MEP, CLP and GMP (**Figure 4F**). We noted the appearance of a population dominantly presented in the *Cnot3-*cKO, which we named KOp (**Figure 4F**). To probe the identity of the KOp cluster, we performed GSEA analysis comparing the gene expression profiles of HSC vs. KOp and found that the KOp cluster exhibited a decreased stem cell gene expression program concurrently with an increased myeloid development program (**Figure S4B**). Pseudotime analysis revealed that the KOp cluster appears to be an intermediate progenitor population between HSC and GMP (**Figure 4G and Figure S4C-D**). The results suggest that while CNOT3 functions in a multilineage manner, its loss initially alters HSC fate decisions more prominently.

**Figure 4.**
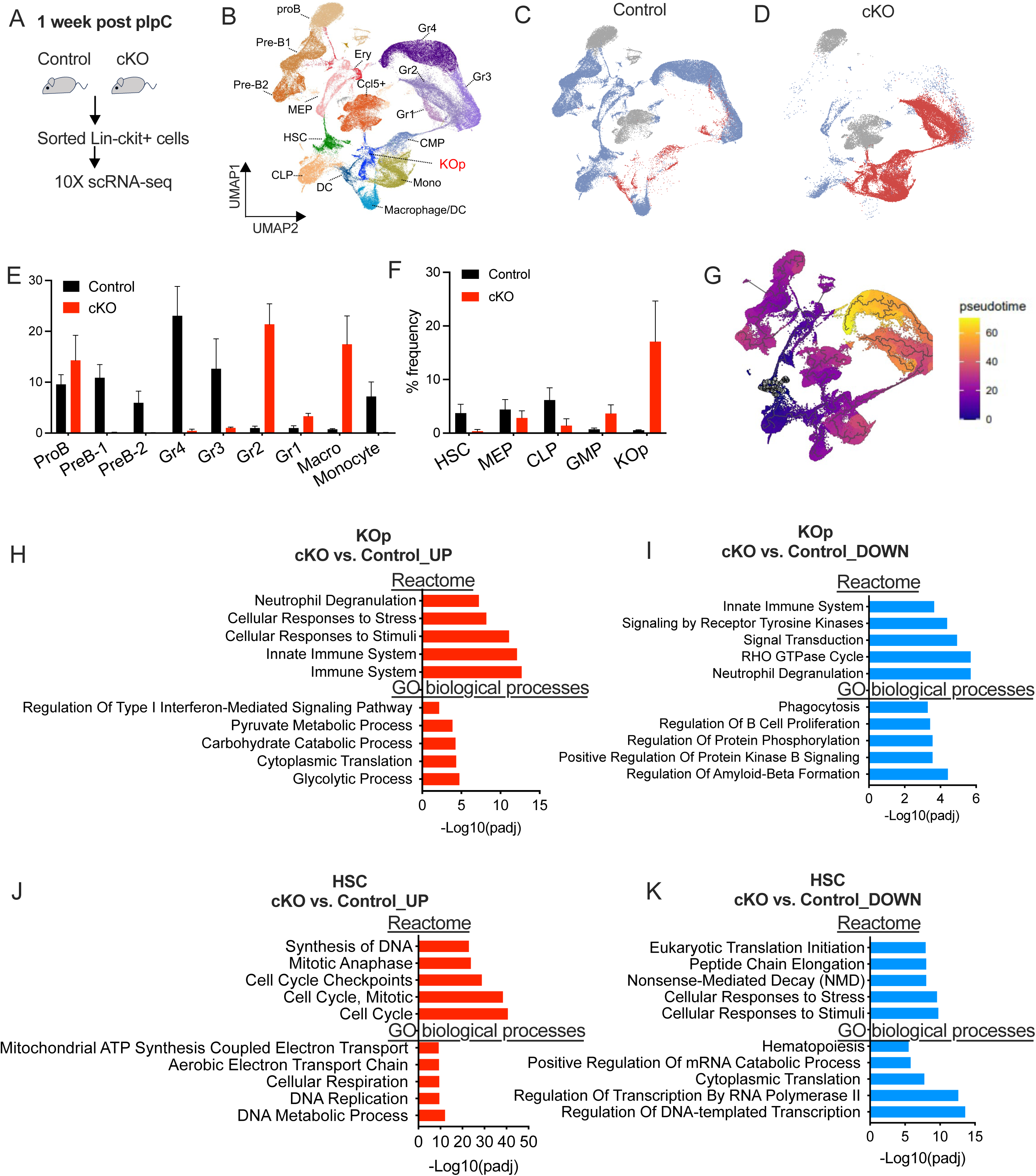
scRNA-seq analysis revealed impairment in multi-lineage development and a defect in HSC compartment upon *Cnot3* deletion. (A) Experimental scheme for single cell (sc)RNA-seq analysis of Lin-ckit+ HSPCs isolated from Control (n=3) vs. cKO (n=3) mice at 1 weeks post-pIpC. (B) UMAP analysis of Single-cell RNA sequencing (scRNA-seq) showing hematopoietic cell populations identified in Control and cKO mice. (C-D) UMAP showing of all hematopoietic clusters of Cnot3 Control (C) vs. cKO (D). (E-F) Frequencies of different populations defined by scRNA-seq analysis in Control vs. cKO. (G) Pseudo-time reconstruction of the hierarchy of cell differentiation by diffusion pseudotime (DPT) algorithm based on scRNA-seq of Control and cKO mice. (H-K) Enrichr analyses of GO biological processes and Reactome enrichment of significant (FDR < 0.05) downregulated and upregulated genes within the KOp population (H and I) and HSC population (J and K) of Control vs.cKO. X-axis: -log10(p value).

To uncover the functional pathways altered in different cell populations upon CNOT3 deletion, we performed differential gene expression analysis in each clusters (**Table S2**) and conducted gene set enrichment analyses using Enrichr (https://maayanlab.cloud/Enrichr/)(26). Within the *Cnot3* Kop population, depletion of CNOT3 resulted in upregulation of genes enriched in immune responses, cellular response to stimuli and several metabolic processes (**Figure 4H**) and downregulation of targets mainly belonging to signal transduction pathways and GTPase cycle (**Figure 4I**). On the other hand, CNOT3 deficient HSCs exhibited strong activation of DNA synthesis and cell cycle (**Figure 4J**) but suppression of genes in mRNA transcription, processing and translation (**Figure 4K**). The results indicated that dysregulation of cell cycle in *Cnot3* cKO HSCs may lead to abnormal cycling and expansion of the aberrant group of stem/progenitor *Cnot3* KOp cells.

### Loss of CNOT3 resulted in increased cell cycling and abnormal division of HSPCs

Given the observed change in the transcriptomes of CNOT3 deficient HSCs, we directly assessed cell cycle status of HSPCs upon CNOT3 depletion. We performed flow cytometry analysis with Ki67 and DAPI staining to examine whether deletion of *Cnot3* impairs HSPCs’ cell cycle progression. We observed a marked reduction in G_0_ quiescent population and an increase in the proportion of cells entering G_1_ and cycling in S/G2/M phase in LSKs and HSCs (**Figure 5A-B and Figure 5C-D**). To further probe if CNOT3 deletion pushed HSPCs to proliferate, we incubated Control vs. cKO LSKs and HSCs with BrdU and tracked the incorporation of BrdU in cycling cells. We noted an increase in percentage of cells in apoptotic sub_G1 in *Cnot3* deficient LSKs and HSCs (**Figure S5A**). Most significantly, we found a large portion of cells were positive for BrdU staining, indicating that CNOT3 deletion drove them into cycling (**Figure 5F-G**). In addition, there was increased DNA content in CNOT3 deleted HSPCs in S phase (**Figure 5H**). These findings indicate that loss of CNOT3 leads to deregulated DNA synthesis and uncontrolled cycling of HSPCs.

**Figure 5.**
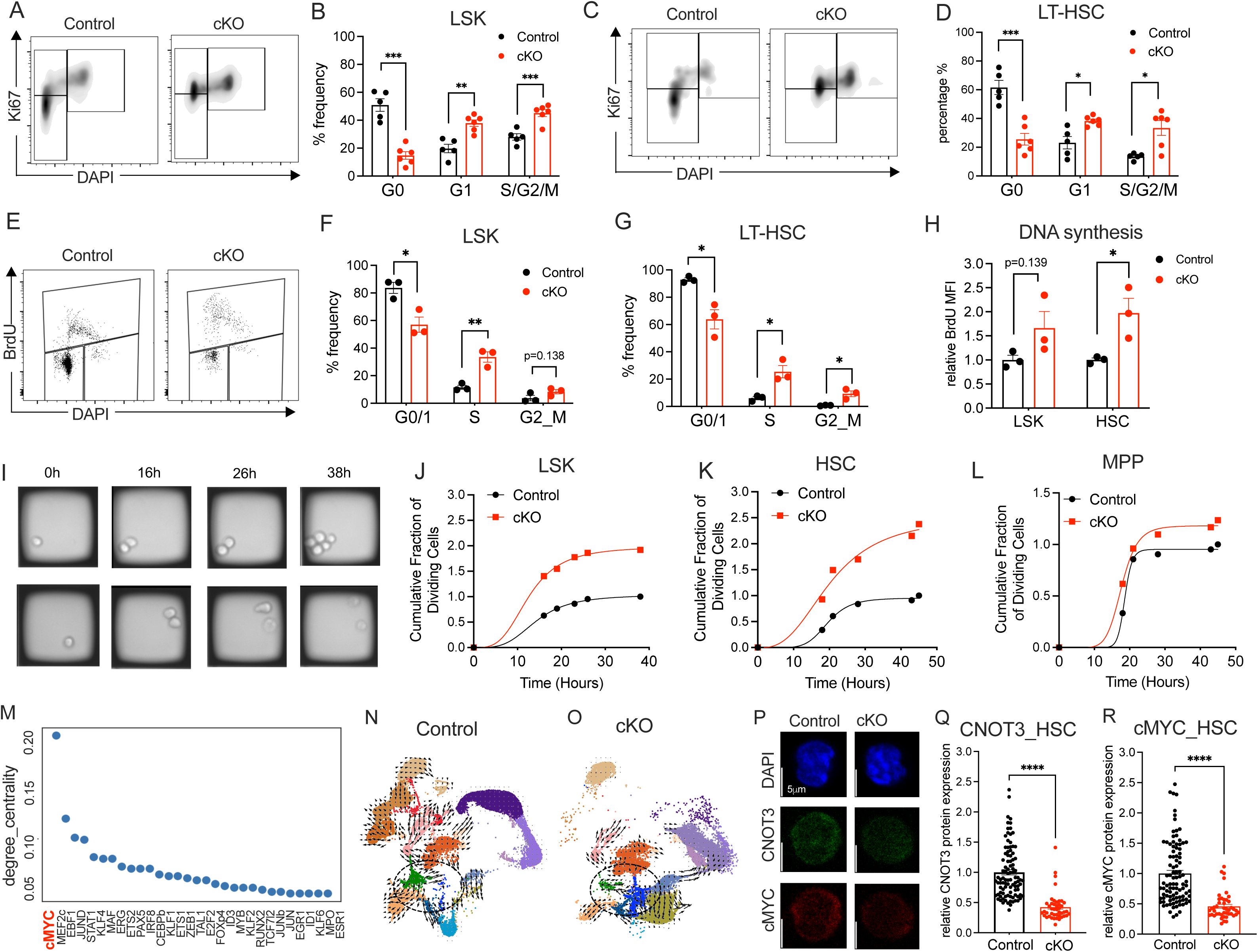
Loss of CNOT3 induced cycling and division of HSPCs. (A) Representative flow plots showing cell cycle assessments by DAPI and Ki67 staining of LSKs from Cnot3 Control vs. cKO mice. (B) Quantitative summary of results in LSK from (A) n=5 for Control and n=6 for cKO. (C) Representative flow plots showing cell cycle assessments by DAPI and Ki67 staining of HSCs (CD150+CD48-LSK) from Cnot3 Control (C) vs. cKO mice. (D) Quantitative summary of results in HSCs from (C) n=5 for Control and n=6 for cKO. (E) Representative flow plots showing cell cycle assessments by DAPI and BrdU staining of LSKs from Cnot3 Control (C) vs. cKO mice. (F) Quantitative summary of results from LSK (E) n=3. (G) Quantitative summary of cell cycle assessments by DAPI and BrdU staining of HSCs n=3. (H) Fold increase in BrdU median fluorescence intensity (MFI) in cells cycling in S phase in Control (C) vs. cKO mice n=3. (I) Representative images of in vitro LSK cell division captured on CellRaft AIR System (Cell Microsystems), shown in brightfield. (J) Cumulative graphs tracking in vitro division of LSKs over the course of 40 h. All data points were normalized to the total division events recorded in Control LSKs, n=3 donors for each genotype. (K) Cumulative graphs tracking in vitro division of MPPs (LSK CD48+ CD150+/-) over the course of 40 h. All data points were normalized to the total division events recorded in Control MPPs, n=3 donors for each genotype. (L) Cumulative graphs tracking in vitro division of HSCs (LSK CD48-CD150+/-) over the course of 40 h. All data points were normalized to the total division events recorded in Control HSCs, n=3 donors for each genotype. (M) Degree centrality scores were used to rank the top 30 transcript factors in driving the differences between HSCs vs. Cnot3P populations. (N-O) Simulation of lineage trajectory upon cMYC loss of function in Control (N) and cKO (O) cells. (P) Representative immunofluorescent (IF) images of IF staining of CNOT3 and cMYC in HSCs. (Q-R) Quantification of normalized IF CNOT3 (Q) and cMYC (R) protein expression in HSCs. (Control n = 96 and cKO n = 44, p < 0.001). (A-R) All graphs show data as mean+/- s.e.m. Two-tailed Student’s t test. ns not significant * p< 0.05, **p<0.01, *** p<0.001.

Next, we wanted to determine whether the unchecked cycling of CNOT3 deficient HSPCs leads to abnormal cell division. We plated single LSKs and HSCs sorted from Control vs. *Cnot3* cKO mice 1 week after pIpC into single wells. We then visualized and tracked cell division in vitro using the CellRaft AIR® System over the period of 2 days (**Figure 5I and S5B**). While there was no notable difference in the median time to first cell division of Control vs. *Cnot3* cKO LSKs or HSCs (LSK CD48-CD150+/-), we observed about two-fold increase in the number of cells completing the first cell division upon CNOT3 depletion (**Figure 5J-K**). Interestingly, the phenotype in MPPs (LSK CD48+CD150+/-) appeared considerably less pronounced (**Figure 5L**). Furthermore, despite more *Cnot3* deficient HSPCs progressed to the 2-cell stage, much fewer cells were found to complete the second division to 4 cells when CNOT3 was deleted (**Figure 5I**). The data suggested that CNOT3 loss resulted in aberrant HSCs’ division, possibly leading to expansion of the stem/progenitor *Cnot3* KOp population.

To further investigate the molecular changes that can be responsible for the dysregulated cell fate and identity in CNOT3 knockout HSCs, we applied CellOracle(27) tool to the scRNA-seq to explore the gene-regulatory networks (GRNs) perturbed upon CNOT3 depletion. We then integrated the GRNs with the pseudotime trajectory to characterize the cell-type-specific GRNs and apply CellOracle simulation to identify known TF regulation of cell identity driving transition from HSC cluster to the *Cnot3* KOp cluster. Degree centrality is used to score the number of genes directly connected to a transcription factor, for which highly connected one is inferred to have biological significance within the tested system. Within our datasets, CellOracle pulled out cMYC as the highest ranked regulator based on degree centrality, along with myeloid related transcription factors including MEF2c, JUND, KLF4 and transcription factors important for specification of B lineages such as EBF1 and IRF8 (**Figure 5M**). Given the critical role of cMYC in maintenance of HSCs function (10, 28, 29), we focused our analysis on cMYC. Simulation of cMYC loss of function showed the strong push to transition from HSCs to *Cnot3* KOp (**Figure 5N-O**), which was aligned with the observed accumulation of functionally defective HSCs (**Figure 3D and 3F-H**). The data suggested that the *Cnot3* cKO HSC phenotype can be at least in part explained by cMYC loss of function. No significant change in *cMYC* mRNA levels when CNOT3 was knockout was noted in both HSCs and *Cnot3* KOp cells (**Table S2**). However, given that we previously found that CNOT3 regulates cMYC translation in leukemia cells(24), we performed immunofluorescent (IF) analyses of cMYC and CNOT3 in sorted HSCs and MPPs. We saw a strong correlation between expression levels of the two proteins (**Figure S5C**). Importantly, we found that efficient depletion of CNOT3 resulted in marked reduction in cMYC protein in HSCs (**Figure 5P-R**). The data strongly suggested that cMYC loss was at least partially responsible for the phenotypes observed in CNOT3 depleted HSCs.

## Discussion

While the role of post-transcriptional regulation has been increasingly implicated in the control of blood stem cells, direct and comprehensive assessments of functional requirement for the regulatory proteins have been limited to a handful of RNA binding proteins. In this study, we interrogated the role of CNOT3, a subunit of the CNOT RNA deadenylation complex, in adult hematopoiesis and in HSPCs. Using a blood specific, murine *Cnot3* conditional knockout model, we showed that the loss of CNOT3 resulted in lineage specific phenotypes i.e., mild alternations in myeloid cells but strong defects in erythroid maturation and abnormal B cells differentiation, which was agreeable with previous study(18). By looking at the changes within the hematopoietic compartments at multiple time points (weeks 1, 2 and 3 post pIpC induction), we captured the most notable phenotype within the HSPC populations where there was first a transient expansion of immunophenotypic HSCs (at week 1 post pIpC induction) followed by the progressive loss of the stem compartments coupled with increase presentation of the multiple potent progenitors (at weeks 2 and 3 post pIpC induction). Importantly, *Cnot3* depleted cells, despite being home to the bone marrow, completely failed to reconstitute in recipient animals. Interestingly, we found that deletion of one allele of *Cnot3*, resulting in 50% reduction of the protein, had no impact on normal hematopoiesis or HSPCs, strongly indicating a dosage dependent requirement for CNOT3 function. The data confirmed out previous observation that shRNA-mediated ablation of CNOT3 to half of the normal level did not impact survival of human HSPCs, further supporting a possible therapeutic window where targeting CNOT3 may have minimal impact on normal cells.

Given the observed expansion of the HSCs at the early time points, we leveraged the scRNA-seq to investigate the transcriptomic profiles of each population upon CNOT3 depletion. Our scRNA-seq revealed the overrepresentation of a “new” population, which we termed KOp in cKO mice. Pseudotime and cell fate analyses characterized KOp population to constitute of direct progenies from HSCs but not committed to differentiation. The observation was somewhat similar to what we found before in *Mettl3* conditional knockout HSCs where there was an emergence of an aberrant intermediate state of HSCs, giving rise to MPP-like populations (30). In addition, using scRNA-seq data, we were able to look into the changes specific for each cell cluster, which further illustrate the differential function of CNOT3 in different cell types. We saw that the depletion of CNOT3 resulted in distinct changes in the transcriptomes of HSCs and KOp populations. While loss of CNOT3 in HSCs led to alternations in DNA replication and cell cycle control, CNOT3 deficient KOp exhibited abnormal metabolism; activated cellular responses to stress and immune pathways.

At the cellular levels, in agreement with the transcriptomic data showing cell cycle dysregulation in *Cnot3* cKO HSPCs, we showed conclusive data demonstrating a marked increase in cells cycling and significant reduction of HSPCs staying within the G0/G1 quiescent state. Depletion of CNOT3 also led to heightened DNA synthesis rate. It has been shown that CNOT3 positively regulates mRNA stability and reduces mRNA level of *Cdkn1a* aka p21 mRNA, which at least in part responsible for the cell cycle defects upon CNOT3 loss of function(31). We also noted upregulation in *Cdkn1a* across clusters in the scRNA-seq dataset, indicating that CNOT3’s regulation of CDKN1A may be part of the mechanisms underlying this phenotype. In addition, we employed a single cell-based assay to track division of HSPCs. Interestingly, while we observed increased in the number of first division in both HSCs and MPPs, the phenotype appeared to be more pervasive in HSCs. Moreover, our analyses of trajectory and regulatory network revealed an involvement of cMYC in controlling cell fate decision of HSCs. cMYC has been shown to be important in many aspects of HSC biology (32, 33) but its exact function in HSPCs depends on cMYC expression level and to some extent context dependent (10, 29). The previous link between CNOT3 and cMYC (24) prompted us to examine cMYC regulation by CNOT3 in this setting. The results indeed suggested a post-transcriptional control of cMYC expression by CNOT3 and that this regulatory connection may be more critical for mediation of HSC division and commitment.

Overall, our data pointed to a more discrete role of CNOT3 in different cell types and the ability to investigate its multifaceted functions within the blood hierarchy was particular informative. Our scRNA-seq dataset unveiled regulation of different biological pathways by CNOT3 within particular cell populations, providing a strong foundation for future studies into the context-dependent control of gene expression programs. Importantly, we established CNOT3 as a critical regulator of cell identity and required for maintenance of stem cell activity in the hematopoietic system.

## Material and methods

### Animal research ethical regulation statement

Protocols were approved by the UBC Animal Care Committee (ACC) and ensured to be in compliance with legislations, policies and procedures under the Canadian Council on Animal Care (CCAC) and Canadian Association for Laboratory Animal Medicine (CALAM) Standards of Veterinary Care. The colonies were kept in alternating 12-hour light and dark cycles at Animal Research Centre (ARC) at BC Cancer Research Institute (BCCRI).

### *Cnot3* hematopoietic conditional knockout model

*Cnot3* flox/flox was obtained from Guang Hu laboratory at National Institute of Environmental Health Sciences, Research Triangle Park. To generate hematopoietic specific *Cnot3* conditional knockout model, *Cnot3* flox/flox mice were crossed with B6.Cg-Tg(Mx1-cre)1Cgn/J (i.e. Mx1 Cre) mice, which were obtained from the Jackson laboratory (Strain#:003556) to generate Cnot3 flox/flox Mx1 Cre+ conditional knockout (cKO) mice. Cnot3 flox/flox and null for Mx1 Cre (Cnot3 flox/flox Mx1 Cre-) mice were used as controls. *Cnot3* deletion in adult mice were induced through exposure to polyinosinic-polycytidylic acid (pIpC) (InvivoGen (HMW) VacciGrade). CNOT3 flox/flox, Mx1-Cre+ or Mx1-Cre-mice between 8 to 12 weeks of age were injected intraperitoneally once a day over two consecutive days with 10 mg/kg pIpC.

### Flow Cytometry and cell sorting

Cell populations in peripheral blood, spleen and bone marrow samples were identified through analysis using multi-parametric flow cytometry. Collected cells were subjected to red blood cell lysis and stained with antibodies at 1:200. Same antibodies were used to sort for LSK cells (Lin-, ckit+, sca+), MPP and HSC (CD48, CD150).

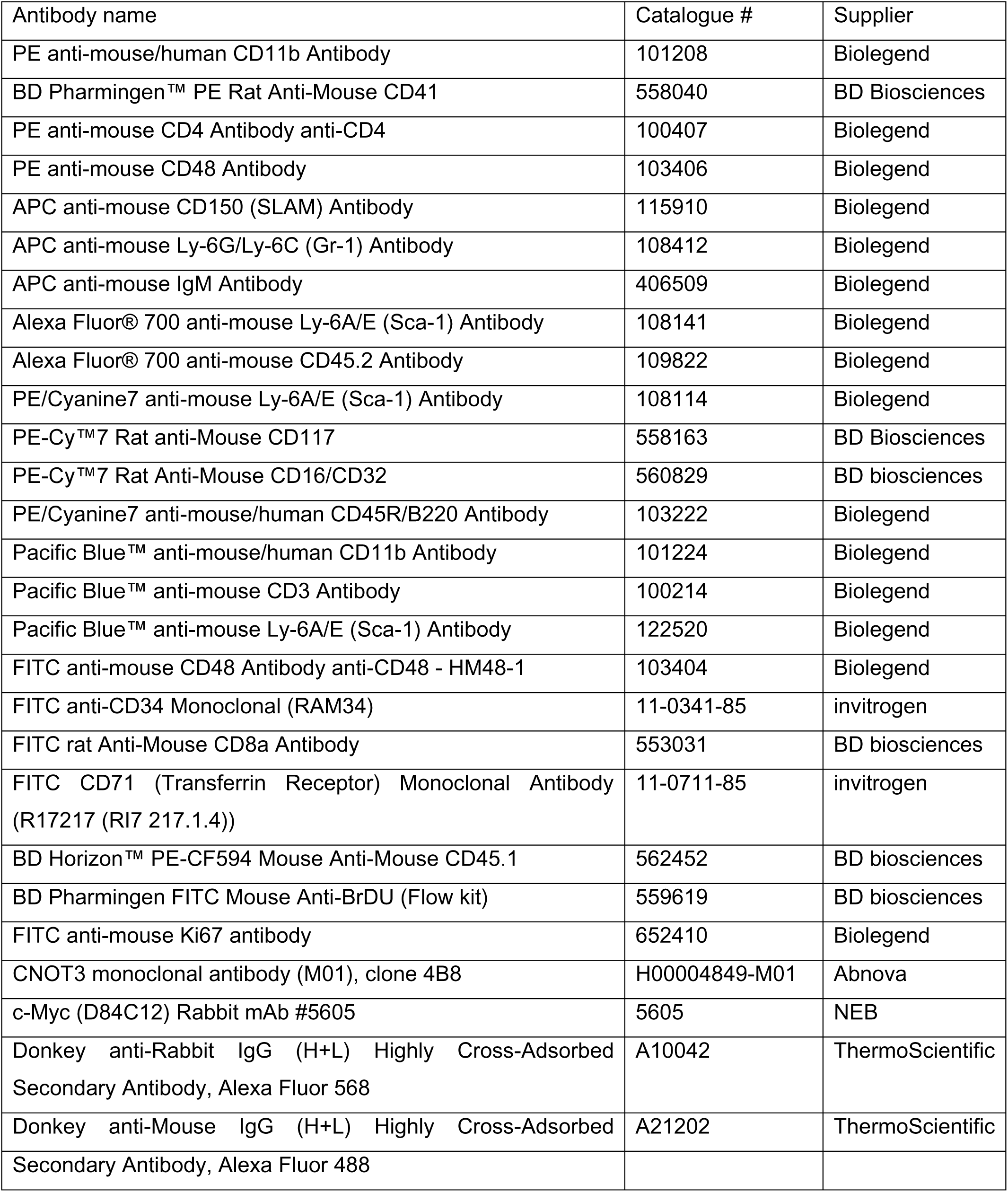

### In vivo Bone Marrow Transplantation Assays

Pep3b female mice expressing CD45.1, congenic to C57BL/6, were used as recipients for transplant assays. Recipients were lethally irradiated with 2 doses of 5 Gy using a cesium irradiator and injected with 2 x 10^6^ total bone marrow cells isolated from the donors. Peripheral blood and bone marrow were then monitored over 24 weeks for CD45.2 chimerism to determine repopulation ability of the donor cells. CD45.1 (recipient) and CD45.2 (donor) cells were detected via flow cytometry.

### ImmunoBlotting

Protein lysates were obtained using 1X Laemmli buffer (Bio-Rad #1610737EDU) and B-Mercaptoethanol. Samples were loaded onto an 4-15% polyacrylamide gel for SDS-PAGE. Proteins were transferred onto a nitrocellulose membrane and blocked for 1 hour (1X TBS-Tween 0.1% + 5% milk). The primary antibodies were diluted to 1:1000 in and left to incubate overnight at 4°C. The antibodies used targeted CNOT3 (mouse, Abnova #H00004849-M01), c-Myc (rabbit, Cell Signaling # #5605) and B-actin conjugated with HRP (mouse, Sigma, # A3854). The secondary anti-mouse or anti-rabbit antibody conjugated with HRP enzyme are diluted to 1:10000. The membranes were covered with a chemiluminescent substrate (HRP Immobilon Western Substrate, Millipore, WBKLS0500) and left to incubate in the dark for 1 minute. Images were taken using Bio-Rad ChemiDocTM System. The values obtained were normalized to the results obtained for actin.

### Cell cycle analysis

Total bone marrow from CNOT3 f/f, Mx1-Cre+ or Mx1-Cre-mice injected with Poly(I:C) were collected and stained with LSK and HSC surface markers then fixed and permeabilized using BD Biosciences Cytofix/CytopermTM buffer (#51-2090KE). Cells were stained with KI67 and DAPI (1:200) and analysed by flow cytometry. BrDU labelling and staining was performed using BD Pharmingen® FITC BrdU Flow kit (#559619). Bone marrows were collected from CNOT3 f/f, Mx1-Cre+ or Mx1-Cre-mice injected with Poly(I:C) (HMW) VacciGrade™ (InVivoGen). BrdU labelling and staining was performed with BD Pharmingen™ FITC BrdU Flow Kit (#559619), as per manufacturer’s instructions.

For BrdU cell cycle analysis, bone marrow cells were resuspended at a concentration of 2 × 106 cells/mL in media (10μM BrdU RPMI + 10% FBS media, supplemented with 10ng/mL each of SCF, IL-3, GM-CSF and IL-6). After incubation for 90 minutes at 37°C, the cells were harvested and stained with LSK and HSC markers. The cells were fixed and permeabilised with BD Cytofix/Cytoperm™ Buffer and BD Cytoperm Permeabilization Buffer Plus. Cells were then treated with Deoxyribonuclease I and stained with FITC-conjugated anti-BrdU antibody (1:25) and DAPI solution (1:100). Flow cytometry was conducted using BD LSRFortessa™ Cell Analyzer.

### Pathological analysis

Formaldehyde fixed organs were sectioned prior to H&E staining. Samples and images were processed and analyzed at ACS Diagnostic & Research Histology Laboratory Services at the Centre for Comparative Medicine at UBC. Slides were reviewed by pathologist.

### Genotyping of Cnot3 and Mx-1 Cre

Ear notches are lysed in 75µL alkaline lysis buffer (NaOH 25mM, disodium EDTA 0.2 mM, pH = 12) heated at 95°C for 2hours. Neutralization reagent is added (40mM Tris HCl) and samples are used for PCR genotyping using Q5® Hot Start High-Fidelity DNA Polymerase. PCR products are run on a 2% agarose gel and imaged on Bio-Rad ChemiDocTM System.

#### Primers and PCR cycles programs

##### CNOT3 flox/flox gene

Forward primer: GAGCTGCCAGCCTCATGTAT Reverse primer: CGGAAGGAAGGGCATGTAGC

PCR program: 95 (3 min)-95 (10s)-57 (1min)-72 (30 s)-Repeat for 34 cycles – 72 (5 min)-4 hold

*Cnot3* deletion:

Forward primer: TGCTTGTGCTCTAGTGTCATCTAGAGTGC

Reverse primer: CTGCCTCCTAGCCTTCTGACCACCA

PCR program: 95 (3 min)-95 (10s) - 57 (1min) - 72 (30 s) - Repeat for 34 cycles – 72 (5min)-4 hold.

MX1-Cre recombinase:

Forward primer: GCGGTCTGGCAGTAAAAACTATC Reverse primer: GTGAAACAGCATTGCTGTCACTT

Internal positive control F primer: CTAGGCCACAGAATTGAAAGATCT Internal positive control R primer: GTAGGTGGAAATTCTAGCATCATCC

PCR program: 94 (1min) - 94 (30s) - 54 (1min)-72 (1min)-Repeat for 34 cycles - 72 (2 min) - 4 hold

### Tracking cell division using the CellRaft® system

Sorted LSKs and HSCs from bone marrows of Control and Cnot3 cKO mice were tracked for cell division on the CellRaft® system from Cell Microsystem. CellRaft® quad arrays 100×100 were coated overnight at 37° with 500µL of laminin-521 (StemCell Cell Adhere® #77003) diluted in PBS (1:10).

After washing once with PBS, quad arrays were seeded with approximately 16,000 LSK cells (Lin-, ckit+, sca+), MPP or HSC (CD48, CD150) in 500 µL BMT media (RPMI,10% FBS, 10ng/mL SCF, 10ng/mL IL-3, 10ng/mL IL-6, 10ng/mL GM-CSF). Arrays were scanned using the CellRaft® AIR system every 4 hours to track cell division. Analysis is performed on the CellRaft® Cytometry software.

### Immunofluorescent (IF) imaging

MPP (Lin-Sca-1+ c-Kit+ CD48+) and HSC (Lin-Sca-1+ c-Kit+ CD48-) cells were sorted and fixed in 10% neutral buffered formalin (NBF) at a concentration of 105 cells/mL for 60 minutes at 37 °C and stored in 70% ethanol.

Slides were prepared by cytospinning 0.2 – 0.8 x 105 cells onto slides at 750 x g for 5 minutes using the Thermofisher Cytospin 4. Cells were permeabilised by immerging slides consecutively in 50%, 70% and 2 x 100% ethanol for 5 minutes each. Cells were blocked with 5% FBS, 0.5% Triton X-100 solution at room temperature for 60 minutes. Cells were incubated overnight at with primary antibodies (Abnova, anti-CNOT3: 1:500 (H00004849-M01) and Cell Signalling Technology, anti-c-Myc: 1:500 (D84C12)), in blocking solution. Cells were washed with 3x PBS-T and incubated with secondary antibodies (Invitrogen, anti-mouse AF488: 1:500 (#A21202), anti-rabbit AF568: 1:400 (#A10042)) in blocking buffer for 60 minutes at room temperature. Following washing with 3x PBS-T, cover slips were mounted using DAPI mounting media (Sigma Duolink ® In Situ Mounting Medium with DAPI #DUO82040). Imaging was performed on the Zeiss LSM800 Confocal Microscope and analysed using the Zeiss Zen ® Software and Fiji ImageJ ®.

### Single-cell RNA-seq analysis

Approximately 10,000 Lin-ckit+ cells were sorted and submitted for single cell RNA-seq analysis using 10X Chromium Single Cell 3’ Gene Expression kit. Samples were sequenced at about 20k reads/cell. Single-cell RNA sequencing (scRNA-seq) data were processed using the CellRanger aligned to the mouse reference genome (mm10). Cell quality control was performed in R (v4.3.1) using the DropletUtils (v1.22.0) and scuttle (v1.12.0) packages. Data normalization and cell cycle regression were conducted using the SCTransform method in Seurat (v5.1.0). Principal Component Analysis (PCA) followed by clustering at a resolution of 0.3, which were manually annotated using established marker genes (9). Pseudotime was performed using diffusion pseudotime (DPT) algorithm within the Scanpy environment. In brief, lineage information was set so that cells were divided into lineage branches or sub-clusters based on known developmental pathways. Root cell was selected for each lineage, representing the starting point of the developmental trajectory. The DPT algorithm was then applied to calculate pseudotime for each cell, mapping its progression along the developmental trajectory. CellOracle (27) was used to predict gene regulatory networks (GRNs) from the single-cell RNA-seq data. GRN predictions were further integrated with pseudotime simulation with GRN predictions and visualized using Plotly in Python.

Reviewer link to single cell RNAseq datasets: https://dataview.ncbi.nlm.nih.gov/object/PRJNA1165832?reviewer=9cmnk5v5c0ugljc7onfthuqeqo

## Acknowledgments

The work was supported by Natural Sciences and Engineering Research Council of Canada (NSERC) Discovery Grant and Canadian Institutes for Health Research (CIHR) Project grant to L.P.V. L.P.V is the Scholar of the American Society of Hematology, Scholar of the V foundation for Cancer Research and is a Tier 2 Canada Research in RNA biology in Hematological Malignancies and is supported by the Michael Smith Health Research Scholar award. L.P.V’s lab is supported by Canadian Institutes for Health Research Project Grant, Natural Sciences and Engineering Research Council of Canada Discovery Grant and the Terry Fox Research Institute New Investigator Award. We thank the staffs of the Animal Research Center at BC Cancer for their technical assistance. We are grateful to our lab members for their discussion and technical assistance.

## Author Contributions

Conceptualization, L.P.V.; Methodology, K.Y., T.T., Q.A.H, M.Y. and L.P.V.; Analysis, K.Y., T.T., Q.A.H, Z.L, E.S. and L.P.V.; Resources, T.T.M.N and L.P.V; Technical support: E.H., N.R., N.N., G.E.; Writing – Original Draft, K.Y., T.T., Q.A.H and L.P.V.; Writing – Review & Editing, K.Y., T.T., Q.A.H and L.P.V; Supervision, L.P.V.; Funding Acquisition, L.P.V.

## Declaration of generative AI and AI-assisted technologies

The authors declare that no generative AI or AI-assisted technologies were utilized in this paper.

---

**Figure S1.**
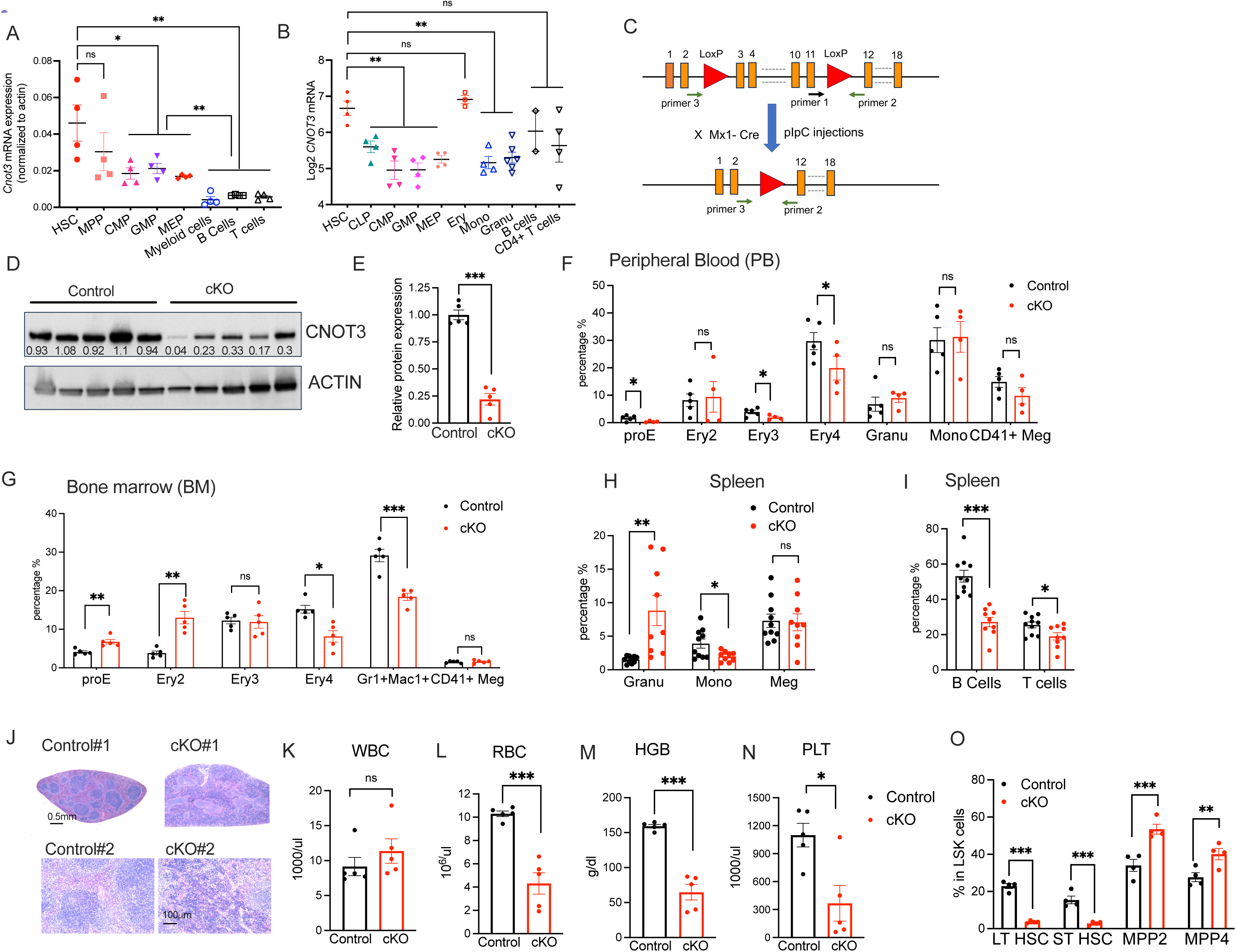
(A) Quantitative RT-PCR measurement of *Cnot3* mRNA expression across hematopoietic stem/progenitor and differentiated cells isolated from mouse bone marrows, n=4. (B) Expression of *Cnot3* mRNA in various mouse blood cell types from publicly available RNA-seq dataset (34) https://www.bloodspot.eu/ (C) Strategy to create hematopoietic *Cnot3* conditional knockout mouse (cKO) and detection of *Cnot3* deletion in the blood system. (D) Immunoblots showing reduction of CNOT3 in cKO upon pIpC induction and *Cnot3* deletion at 3 weeks post-pIpC. ACTIN serves as loading control. (E) Quantitative summary of immunoblot results in (D). (F) Frequencies of multiple differentiate cells in peripheral bloods of Control vs. cKO mice. ProE: Proerythroblasts; Ery2-4: Erythroid 2-4; Granu: Granulocytes; Mono: Monocytes at 3 weeks post-pIpC. (G) Frequencies of multiple differentiate cells in bone marrows of Control vs. cKO mice. Meg: Megakaryocytes at 3 weeks post-pIpC. (H-I) Frequencies of differentiate cells in Spleen of Control vs. cKO mice at 3 weeks post-pIpC. (J) Representative images of H&E staining of spleen samples from Control vs. cKO mice at 3 weeks post-pIpC. (K-N) Analysis of blood counts: (K) White blood count (WBC); (L) Red Blood count (RBC); (M) Hemoglobin (HGB) and (N) Platelets (PLT) of Control vs. cKO mice at the time of death. (O) Frequencies of LT-HSC, ST-HSC, MPP2 and MMP4 in Control vs. cKO mice at the time of death. All graphs show data as mean+/- s.e.m. Two-tailed Student’s t test. ns no significant * p< 0.05, **p<0.01, *** p<0.001.

**Figure S2.**
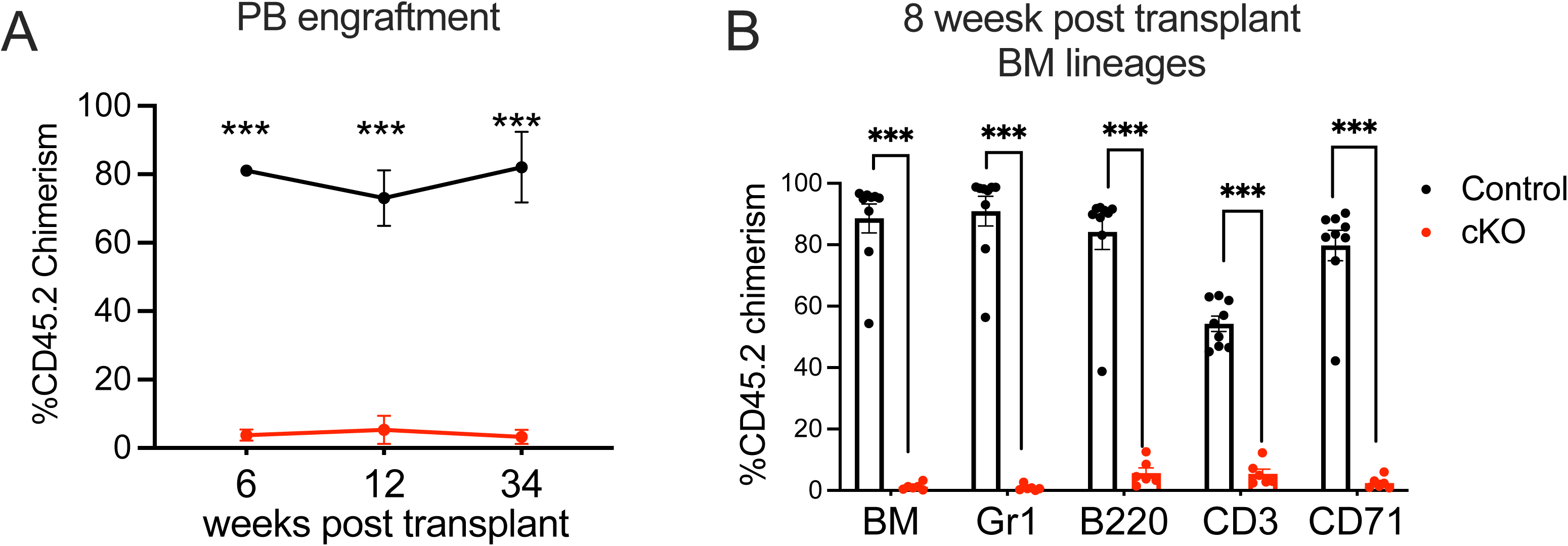
(A) Donor chimerism of Control vs. cKO in peripheral blood (PB) across sequential time points: weeks – 6, 12 and 34 post-transplant. (B) Donor chimerism of Control vs. cKO within various lineages in BM at 8 weeks post-transplant. All graphs show data as mean+/- s.e.m. Two-tailed Student’s t test. ns no significant * p< 0.05, **p<0.01, *** p<0.001.

**Figure S3.**
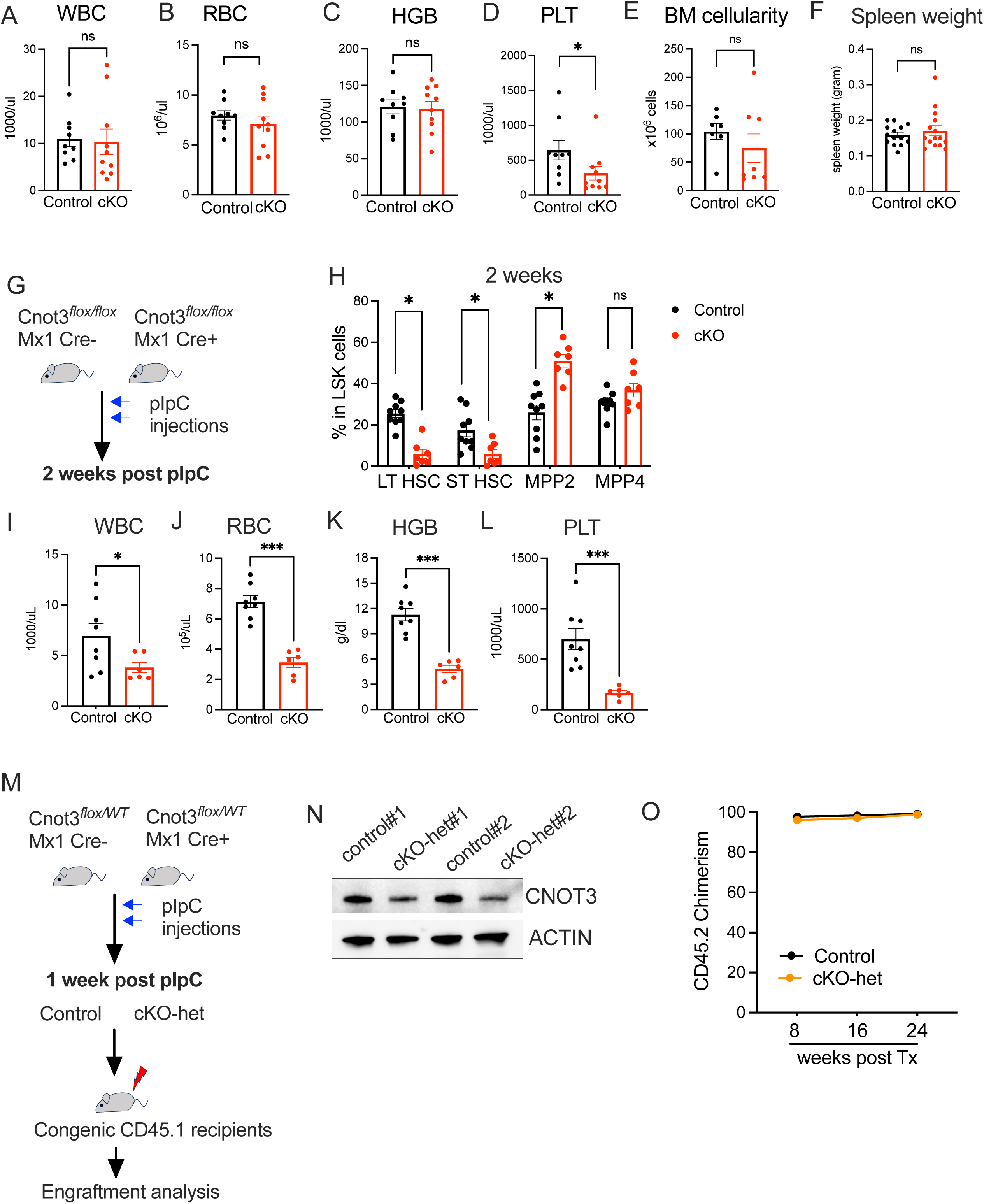
(A-D) Analysis of blood counts: (A) White blood count (WBC); (B) Red Blood count (RBC); (C) Hemoglobin (HGB) and (D) Platelets (PLT) of Control vs. cKO mice at 1 weeks post-pIpC. (E) Total bone marrow cellularity of Control vs. cKO mice. (F) Spleen weight of Control vs. cKO mice. (G) Experimental scheme for pIpC injection and analysis of hematopoietic system upon deletion of *Cnot3* at 2 weeks post-pIpC. (H) Frequencies of LT-HSC, ST-HSC, MPP2 and MMP4 in Control vs. cKO mice. (I-L) Analysis of blood counts: (I) White blood count (WBC); (J) Red Blood count (RBC); (K) Hemoglobin (HGB) and (L) Platelets (PLT) of Control vs. cKO mice at 2 weeks post-pIpC. (M) Experimental scheme for pIpC injection and analysis of hematopoietic system upon deletion of one *Cnot3* allele (cKO-het) at 1 weeks post-pIpC. (N) Immunoblots showing reduction of CNOT3 in cKO-het upon pIpC induction and one *Cnot3* allele deletion at 1 weeks post-pIpC. ACTIN serves as loading control. (O) Donor chimerism of Control vs. cKO het in Bone marrow (BM) across sequential time points: weeks – 8, 16 and 24 post-transplants. All graphs show data as mean+/- s.e.m. Two-tailed Student’s t test. ns no significant * p< 0.05, **p<0.01, *** p<0.001.

**Figure S4.**
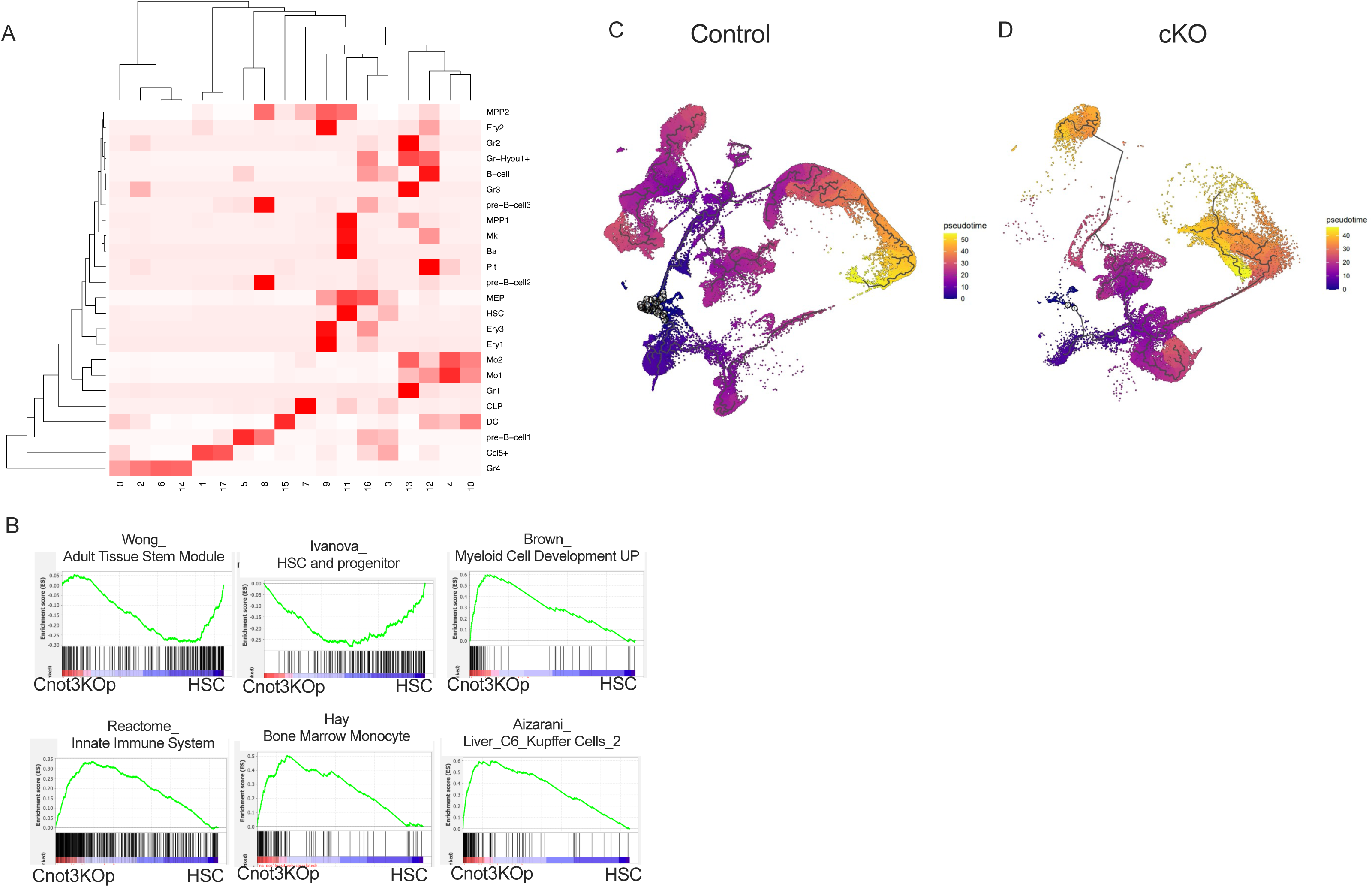
(A) Heatmap showing relative gene expression across all cell types of all marker genes used in UMAP clusters to generate hematopoietic clusters depicted in Figure 4B. (B) GSEA analysis of transcriptomes of Cnot3 KOp vs. HSC populations, showing increased differentiation and immune response signatures in KOp vs. HSC. (C-D) Pseudo-time reconstruction of the hierarchy of cell differentiation by diffusion pseudotime (DPT) algorithm based on scRNA-seq analysis of (B) Control and (C) cKO mice.

**Figure S5.**
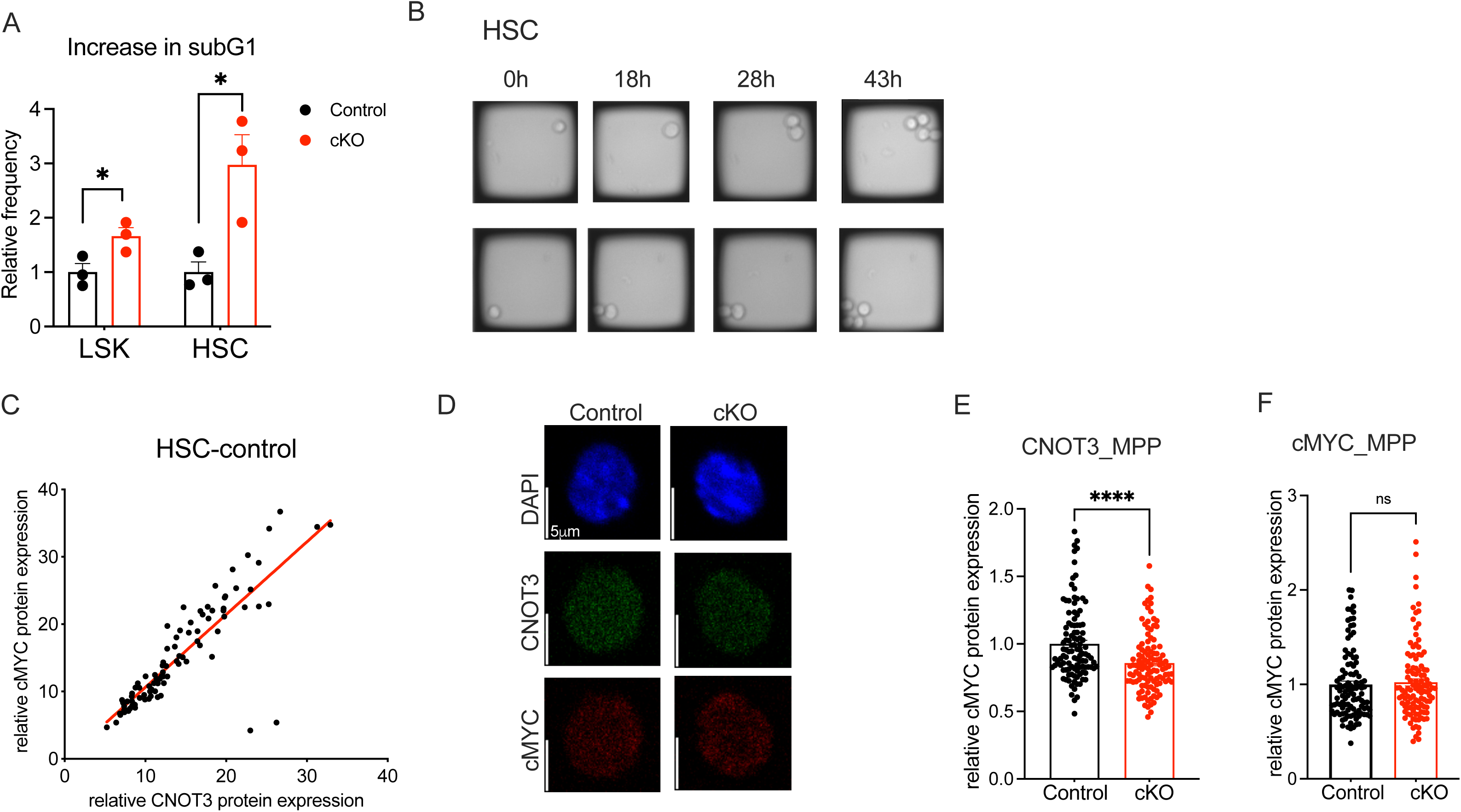
(A) Frequencies of sub-G1 populations within LSK and HSC in Control vs. cKO mice, n=3. All graphs show data as mean+/- s.e.m. Two-tailed Student’s t test * p< 0.05. (B) Representative images of in vitro HSC cell division captured on CellRaft AIR System (Cell Microsystems), shown in brightfield. (C) Correlative analysis of CNOT3 and cMYC protein expression in HSCs. p< 0.0001. (D) Representative immunofluorescent (IF) images of F staining of CNOT3 and cMYC in MPPs. (E-F) Quantification of normalized IF CNOT3 (Q) and cMYC (R) protein expression in MPPs. (Control n = 108 and cKO n = 111, p < 0.001).

